# VARIABILITY IN THERMAL SENSITIVITY OF ADULT REPRODUCTIVE DIAPAUSE IN THE SPRUCE BEETLE, DENDROCTONUS RUFIPENNIS

**DOI:** 10.1101/2023.12.11.570947

**Authors:** Marianne E. Davenport, Barbara J. Bentz, E. Matthew Hansen, Gregory J. Ragland

**Affiliations:** Department of Integrative Biology, University of Colorado Denver, 1151 Arapahoe St., Denver, CO 80204, U.S.A; Rocky Mountain Research Station, USDA Forest Service, 860 North 1200 East, Logan UT 84321

**Keywords:** overwintering, life cycle, bark beetle

## Abstract

Diapause induction and termination, which regulate seasonal insect life cycles, can be highly variable within and among populations because these traits are often genetically variable and sensitive to environmental fluctuations. Both types of variation may influence how insect populations respond plastically or evolutionarily to changing climates. In this study we assessed variability in the expression of reproductive diapause in spruce beetles (*Dendroctonus rufipennis*), a major forest pest whose life cycle timing is regulated by both a prepupal and adult reproductive diapause. Prepupal diapause is facultative and dependent on temperature in the field, whereas previous studies suggest that adult reproductive diapause is effectively obligate. We tested for variability in adult reproductive diapause termination within and between two populations of spruce bark beetle collected from sites in Colorado and Wyoming and reared under warm, summer-like conditions in the laboratory. We also sampled beetles from under tree bark during the fall and spring to estimate how reproductive diapause might terminate naturally in the field. We present evidence that though most beetles induce and do not terminate diapause under constant warm conditions, a small proportion of females from both populations developed mature ovaries and successfully reproduced under warm conditions in the lab. Previous studies have suggested that most beetles require exposure to relatively low temperatures for several weeks to months to terminate diapause in the lab. We found that most female beetles sampled in the field had mature ovaries relatively early in the fall, suggesting that exposure to transiently low temperatures in the field may rapidly terminate adult reproductive diapause. Thus, adult reproductive arrest may primarily act as a block to prevent offspring production prior to winter and appears unnecessary for survival overwinter. Overall, our data do not suggest that major shifts in spruce beetle life cycles as mediated by adult reproductive diapause are immediately imminent with changing climates but, if the variability that we observed is heritable, adult reproductive diapause may have some capacity to evolve in both populations.

**\*\* Note \*\*** all figures/tables designated with ‘S’ are supplemental figures/tables that will appear only in the supplement in the published version. They are provided here alongside the main figures/tables, in-line in the order that they appear in the text for convenience.

## Introduction

Seasonal environments impose strong natural selection to synchronize insect development and reproduction with benign, resource-rich conditions, while mitigating exposure to harsh or resource-poor conditions. Diapause is a type of insect dormancy and is a common adaptive strategy facilitating synchronization via regulation of the timing of diapause entry and exit. Diapause may occur in any life history stage and is typically characterized by reduced metabolism, suppressed development, and increased stress hardiness (Denlinger, 2022).

Diapause induction and termination are cued by environmental variables such as photoperiod, thermoperiod, and resource scarcity (Tauber *et al*., 1986). Moreover, diapause timing is often variable within and among populations (Masaki, 1961, 2002; Ragland *et al*., 2019). Thus, changes in environmental conditions can cause both plastic and evolutionary changes in seasonal timing (Bradshaw & Holzapfel, 2008; Tobin *et al*., 2008).

Many studies document how recent changes in climate are altering the timing of life history events within a season, which in turn may affect trophic interactions and range expansion (Visser & Both, 2005; Altermatt, 2010; Forrest & Thomson, 2011; Buckley *et al*., 2015; Bentz *et al*., 2019). Longer and warmer growing seasons associated with climate change can cause faster development, which can also allow populations to complete more generations in a year (increased voltinism; Jacques *et al*., 2019)), assuming temperatures do not exceed the threshold for optimal development (Deutsch *et al*., 2008). However, there are alternative scenarios where longer growing seasons simply extend the duration of certain life cycle stages, or where interactions between thermal sensitivity of development rate and photoperiodic diapause induction could actually reduce voltinism (Forrest, 2016).

In addition to effects on developmental rates, changes in temperature can alter the timing of diapause entry or exit. In particular, exposure to relatively warm temperatures during a sensitive life stage can prevent facultative diapause induction (Dyer & Hall, 1977; Nunes & Saunders, 1989), even in insects where diapause appears to be obligate under typical, cooler temperatures in the field (Dambroski & Feder, 2007). This can result in additional generations if conditions are permissive, but could also result in high mortality if the environment does not allow growth, development, and/or reproduction necessary to attain a winter-hardy life stage later in the life cycle (Dambroski and Feder 2017). The consequences of averted diapause may be particularly difficult to predict in insects capable of overwintering in multiple life stages. For example, some species of bark beetles have life cycles that vary between semivoltine and univoltine across geography, depending on the environmental sensitivity of two or more possible overwintering stages (Schebeck *et al*., 2017). Even in species where there is one primary overwintering stage, geographic variation in environmental sensitivity may render different populations effectively obligate diapausers (univoltine) or facultative diapausers that may have multiple generations per year when conditions are permissive (Schebeck *et al*., 2022).

The spruce beetle (*Dendroctonus rufipennis* Kirby, Coleoptera: Curculionidae, Scolytinae) is a native disturbance agent of North American spruce (*Picea* spp.) with a flexible diapause/overwintering strategy that may be particularly sensitive to warming climates. It occurs throughout the range of its host trees across western North America from sea level at northerly latitudes (e.g., Alaska) to over 3,600 m in the southern U.S. Rocky Mountains (e.g., northern New Mexico; Holsten, E.H., 1999). Generation time can vary from 1-3 years across this range, where populations experiencing warmer mean temperatures generally have shorter generation time (Knight, 1961; McCambridge & Knight, 1972). Spruce beetle voltinism is regulated by winter diapause that can occur in two life stages; a facultative diapause in the prepupal stage, or final larval instar (Dyer & Hall, 1977; Hansen *et al*., 2001, 2011) and a reproductive diapause in adult beetles that typically requires some exposure to cold temperatures to terminate (**Fig. 1**) (Massey & Wygant, 1954; Safranyik *et al*., 1990; Bleiker & Willsey, 2020).

**Figure 1:**
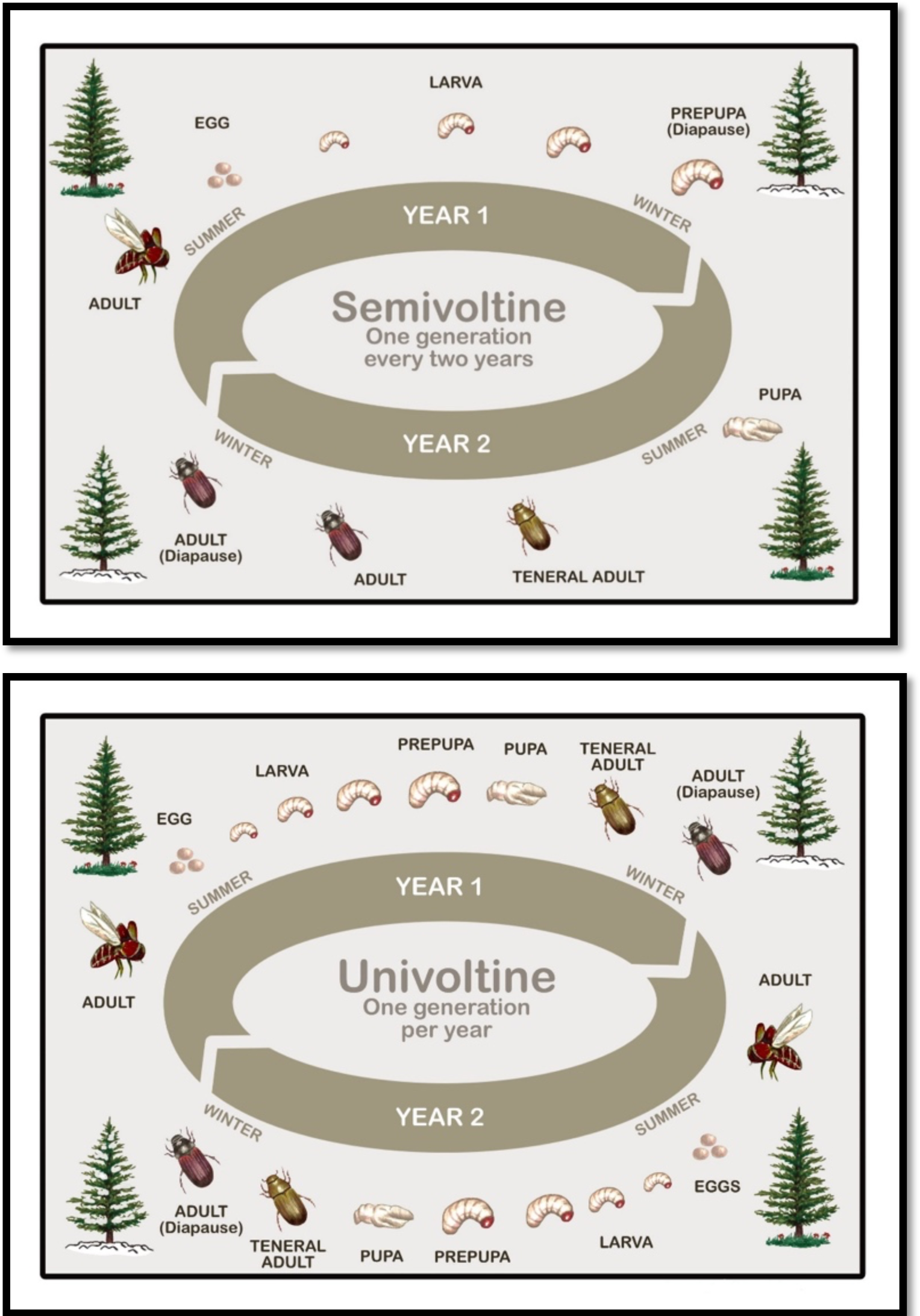
Typical semivoltine (top) and univoltine (bottom) life cycles of *Dendroctonus rufipennis*. Semivoltine beetles overwinter twice, once as a prepupa, and once as an adult, whereas univoltine beetles develop directly to an adult overwintering stage by averting prepupal diapause.

Spruce beetle facultative prepupal diapause is relatively well studied and is induced by temperatures consistently below 15°C in populations sampled from sites ranging from British Columbia south through Colorado and Utah (Dyer & Hall, 1977; Hansen *et al*., 2001, 2011). Consistent exposure to temperatures above 15°C during late summer will avert the facultative prepupal diapause with development progressing to the adult stage prior to winter (Hansen *et al*., 2011). Adults emerge the following summer resulting in a univoltine (one generation per year) life cycle. If prepupal diapause is induced by relatively cool temperatures, the first winter is spent as a prepupa and a second winter is typically spent as an adult leading to a semivoltine lifecycle (one generation every two years). In the less common three-year life cycle, the first winter is passed as an immature larva, the second as a prepupa, and the third as a brood adult.

Research has been comparatively limited on adult reproductive diapause development, but there are multiple lines of evidence suggesting that it is not obligate in the sense that some individuals may become reproductively capable without overwintering. An early natural history study by Massey and Wygant (1954) compared adults that had overwintered as prepupae and then developed to adulthood (i.e., no chilling as an adult) with hibernating adults that had overwintered in the host tree (i.e., experienced chilling as an adult). Although the majority of adult beetles that had not overwintered did not bore into fresh phloem to initiate egg galleries, a few did, suggesting that at least some proportion of adults were reproductively viable without needing overwintering. Likewise, Schebeck et al. (2017) notes unpublished observations of some spruce beetle male-female pairs from a southern location producing eggs without cold exposure. Finally, Bleiker and Willsey (2020) conducted a large, controlled experiment showing that exposing adult spruce beetles to cold temperatures followed by warm temperatures greatly reduced time to emergence from infested logs. However, some adult beetles did emerge from infested logs without any simulated winter cooling, with an even smaller, but non-zero proportion emerging without a pronounced delay. Interpreting these observations is somewhat complicated because adults that have developed from overwintered prepupae will often emerge from infested trees in the fall, relocate to a lower position near the base of the tree, then overwinter as an adult (Massey & Wygant, 1954). Thus, previous results suggest that some proportion of adult spruce beetles could be reproductively mature and produce offspring without winter chilling. There is precedent for this type of variability in the closely related species, *Dendroctonus simplex*; a portion of non-overwintered beetles from a population at the southernmost edge of the species range were found to avert the obligate diapause observed in more northern populations (McKee & Aukema, 2015).

Flexibility of adult reproductive diapause in spruce beetles has important implications for population dynamics with changing climates. The univoltine life cycle for this species occurs when favorable late summer temperatures fail to induce prepupal diapause, allowing for pupation and adult eclosion before the onset of winter (Hansen *et al*., 2001, 2011). Consecutive warm summers have been correlated with regionalized outbreaks (Berg *et al*., 2006), likely due to the exponential increases in population growth rate observed when populations transition from semivoltine to univoltine (Hansen & Bentz, 2003; Werner *et al*., 2006). If adult reproductive diapause can also be averted without obligatory chilling, that could provide additional routes to changes in voltinism. Warming temperatures, however, could also result in fractional or asynchronous voltinism as seasonality and diapause cues are disrupted (Van Dyck et al. 2015; Bentz et al. 2022). Even if aversion of adult diapause without chilling is relatively rare, evolutionary responses to warming climates could shift lifecycle timing if variation within populations is heritable.

We used a combination of laboratory manipulations and field observations to investigate the flexibility of adult reproductive diapause in spruce beetles. Diapause-related traits are often highly variable across geography in many insects including bark beetles (Masaki, 1961; Schebeck *et al*., 2017). We thus conducted laboratory experiments using spruce beetles collected at two geographic sites varying by more than 4 degrees latitude to test if adults could become reproductively mature without exposure to cold temperatures (i.e., no reproductive diapause induction), and if the proportion of non-diapausing individuals was higher in a southern vs northern population. We also measured ovarian maturity from field sampled beetles to determine adult reproductive diapause status prior to and following winter.

## Materials and Methods

### Field Collection of Infested and Green Trees

We harvested spruce beetle-infested Engelmann spruce (*Picea engelmannii* Parry ex Engelmann) from a southern location (Guanella Pass, Colorado) and a northern location (Togwotee Pass, Wyoming) (**Fig. 2**). The locations were chosen based upon data from the USDA Forest Service Aerial Detection Survey (ADS) and 2017 ground surveys showing that both areas had high levels of beetle activity. In June and July 2018, we cut one Engelmann spruce at each site that had been infested the previous year by spruce beetles (**Table S1**).

**Figure 2:**
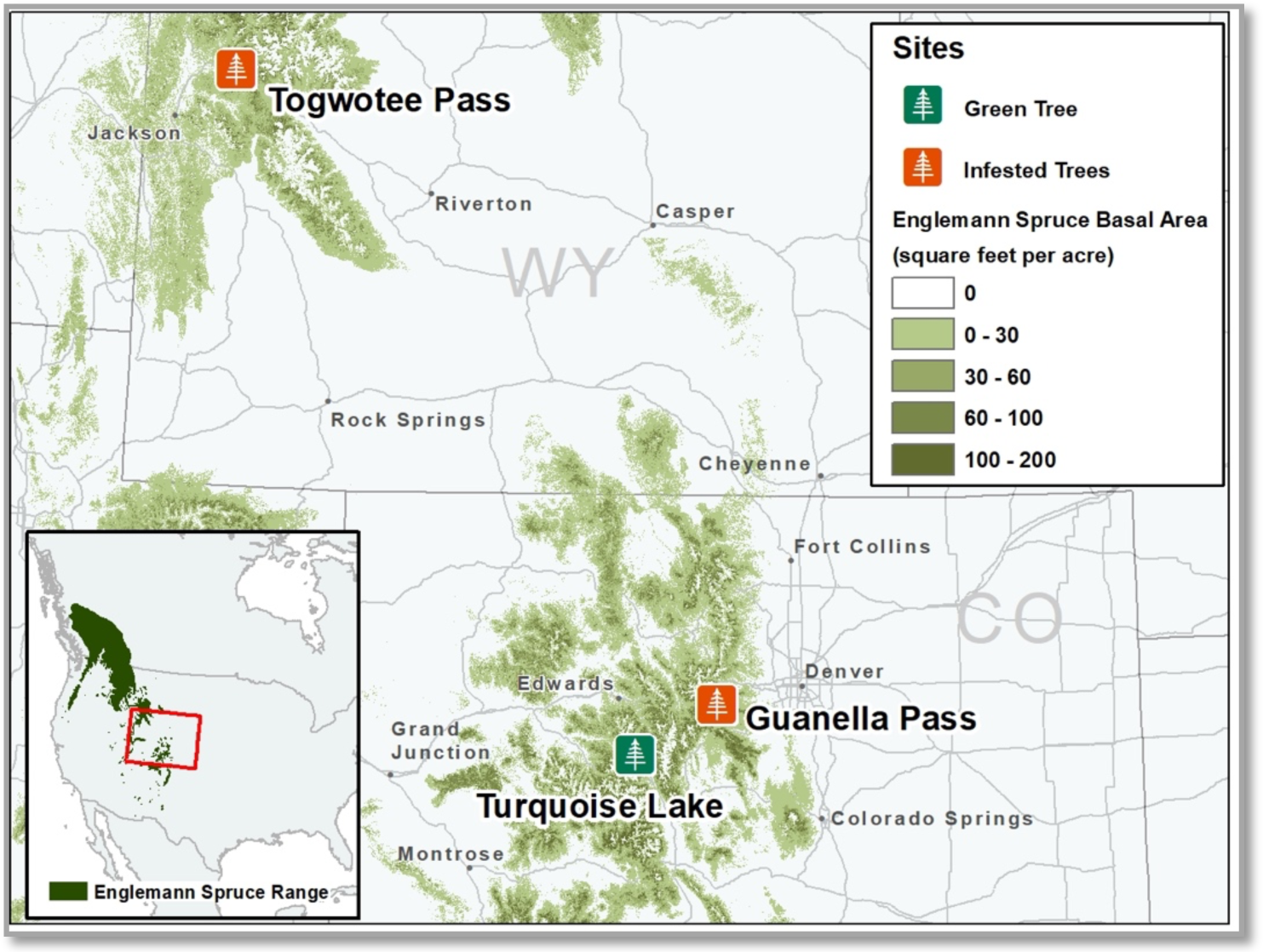
Spruce beetle-infested Engelmann spruce were harvested from a northern population in Wyoming and southern population in Colorado (orange icons). An un-infested Engelmann spruce was harvested from a site in Colorado (green icon), and was used for mating experiments. Distribution of Engelmann spruce in North American shown in inset (data from https://databasin.org/datasets/8f64f9f5e793419eb38ca15580a29d7c/), and Engelmann spruce basal area is shown in monochromatic green (data from https://www.arcgis.com/home/item.html?id=4ebf103ddeeb4766a72e58cb786d3ee2).

Cut bolts from both sites contained a mixture of prepupae, pupae, and adults newly eclosed after prepupal overwintering, suggesting a semivoltine lifecycle (Hansen *et al*., 2001). On arrival in the laboratory, infested bolts were immediately stored in temperature-controlled rooms at 22°C ± 1°C. Each bolt was placed in an individual mesh emergence enclosure to monitor emergence. All adults emerging from these bolts had not experienced cold and were used to assess if adult reproductive maturity could be attained without experiencing cold temperatures (**Fig. 3**).

**Figure 3:**
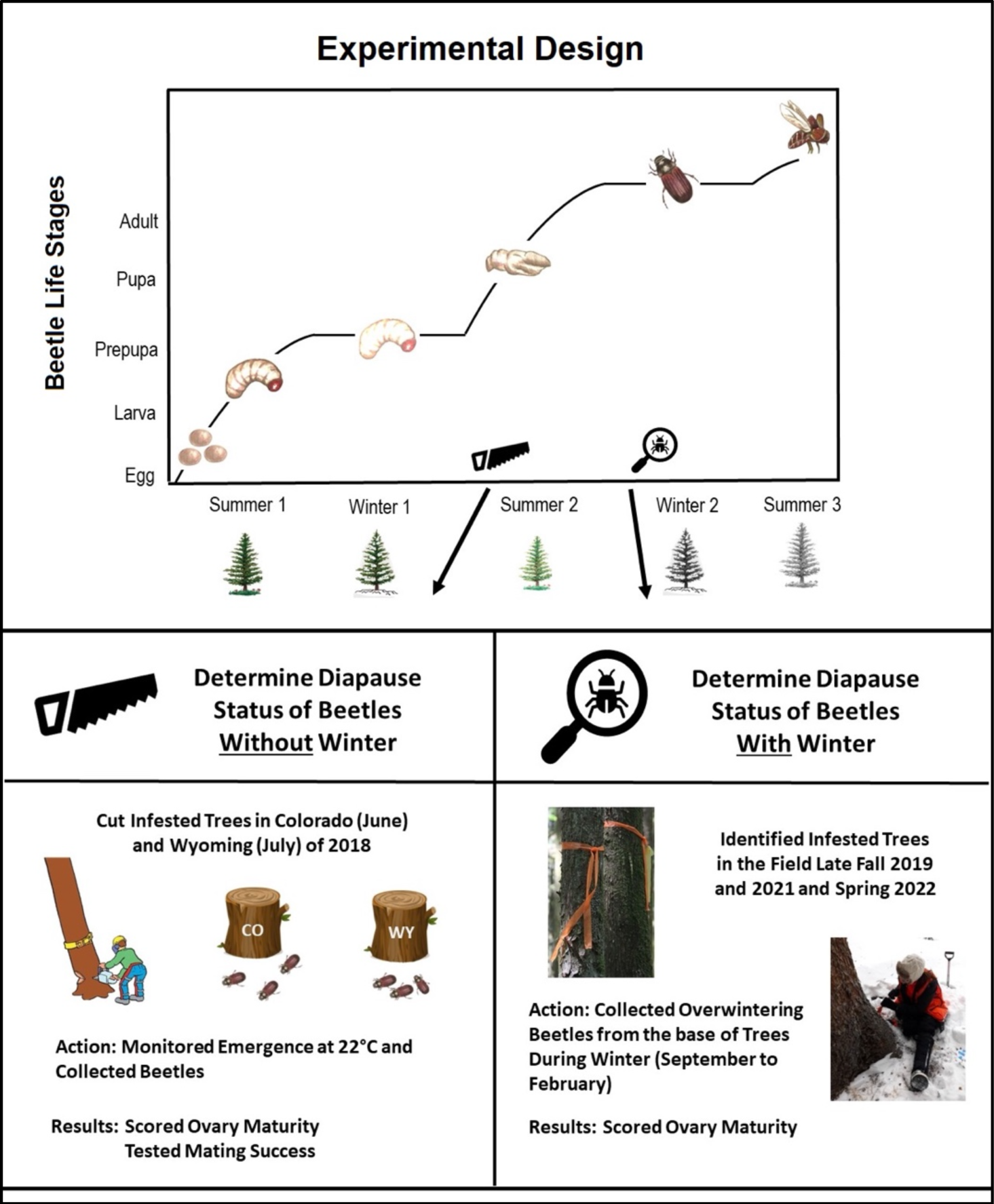
Conceptual experimental design. *Top*: Semivoltine spruce beetle life cycle. *Bottom left*: 2018 experiment details that correspond to life cycle figure with a saw icon. We cut infested trees from Colorado and Wyoming after the beetles completed prepupal diapause. *Bottom right:* 2019 experiment details that correspond to the life cycle figure with a magnifying glass life cycle. We field-collected beetles from Colorado during the winter while they were adults.

To provide bolts for beetle mating trials, we harvested one green Engelmann spruce near Turquoise Lake on the San Isabel National Forest in August 2018. We selected the tree from a stand with no evidence of spruce beetle activity based on ADS data and ground surveys (**Fig. 2**). After felling, 15 bolts approximately 30-35 cm in length, were cut and transported to the University of Colorado Denver. The ends of all bolts were sealed with paraffin wax to reduce desiccation and stored at 4°C ± 1°C.

Data from the closest weather stations to each site were used to compare historical (1999 – 2019) and contemporary thermal conditions. The Colorado site had relatively warmer monthly mean temperatures on average (**Fig. S1**) and more annual degree days > 3°C (929 ± 64) compared to the Wyoming site (789 ± 92).

### Diapause status of laboratory-reared beetles

We monitored adult emergence from infested bolts from both sites, collecting beetles every other day from the start of the experiment (day 0; when bolts were placed at 22°C) until five days passed with no newly emerged beetles (day 246). Beetles from Colorado started emerging after 25 days, while Wyoming-origin beetles started emerging immediately. However, after a pulse of beetle emergence (n=188) on day one, no more beetles emerged from the Wyoming-origin bolts until day 25. This initial pulse of emerged adults was likely re-emerged parent beetles that overwintered following oviposition (Hansen & Bentz, 2003). No beetles emerging prior to day 25 were used for any subsequent experiments or observations. We separated emerging adults by sex (Lyon, 1958). Only adults that emerged from infested bolts from days 50 to 116 were used in mating experiments (N= 180 total beetles, see below). These beetles were either used immediately or stored in petri dishes at 4°C with moist filter paper until use (mean of 4 days of storage). All remaining beetles throughout the emergence period were immediately frozen.

Adult diapause in spruce beetle has been characterized as a lack of reproductive competence (Safranyik *et al*., 1990) implying that adults arrest development prior to ovary maturation. To assess diapause status, we scored both ovary maturity and reproductive success of adults emerging from infested bolts.

We assessed ovary maturity by dissecting 83 Colorado and 94 Wyoming female beetles that emerged from infested bolts. Beetles were randomly sampled across the emergence distribution after day 100 such that sample sizes were approximately proportional to the number of adults that emerged at each time point (see example for CO beetles in **Fig. S2**).

We only sampled after 100 days because prior to this the interval between samples was long enough that some beetles were frozen after dying, compromising internal tissue morphology. All samples collected at 100 days or later for ovary dissection were confirmed alive at the time of freezing. We assessed whether females emerging in the warm lab conditions had mature ovaries by evaluating reproductive organ development based on size and quality of ovarioles. Live-frozen female beetles were removed from the freezer and placed onto a dissecting dish with phosphate-buffered saline (PBS). Elytra, wings and heads were removed, openings were made in the metathorax and ventral sternites, and ovaries were removed.

Determination of development status followed Ryan (1959). We scored ovaries as undeveloped if no oocytes were present in the ovarioles (**Fig. 4a**) and developed if oocytes were present in at least one ovariole (**Fig. 4b**).

**Figure 4:**
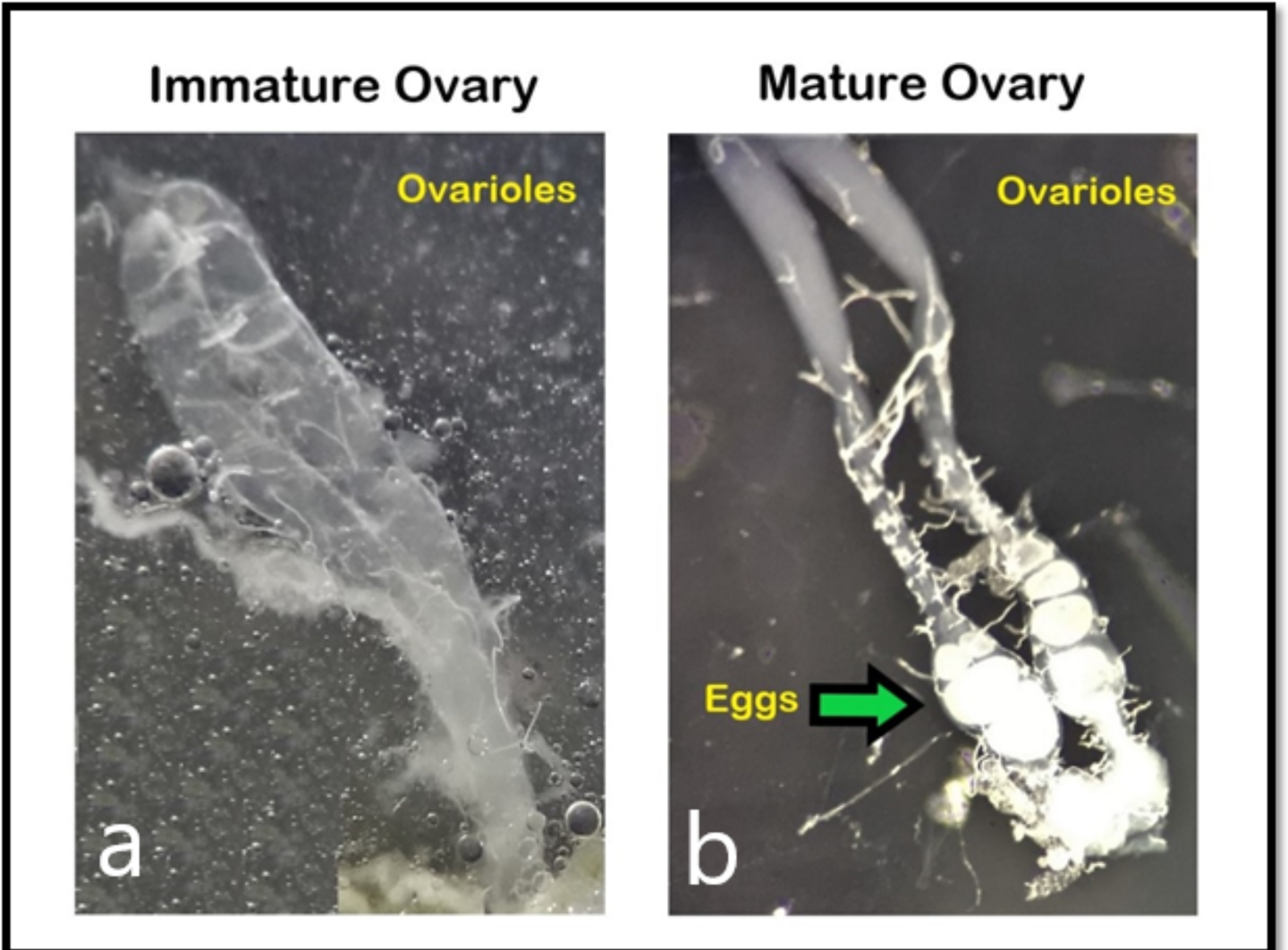
Microscopic images of immature (a) and mature (b) spruce beetle ovaries. 2 ovarioles (of 4 total) are shown for each beetle.

Presence of oocytes in the ovaries does not guarantee that female beetles can successfully mate. Therefore, we also measured reproductive success of 45 male-female pairs of beetles from both sites that emerged between 54 and 116 days after being placed at 22°C. We inserted male-female pairs into holes drilled into the phloem on the cut-side of green, uninfested bolts spaced at 6.35 – 8.89 cm. We first inserted a female into the starter hole, followed by a male after the female disappeared into the phloem. If a male hesitated to join the female, he was replaced with another male. Holes were covered with mesh screen, the bolts inverted and fully enclosed in mesh screen, and placed at 22°C. Sixty days after a bolt was manually infested, it was moved to an incubator set at 4°C to halt development. Cold storage prevented continued beetle movement into galleries from other mating pairs and slowed fungal growth (Eidson *et al*., 2018). After 6 to 76 additional days we carefully removed the bark from each bolt, keeping galleries intact, then counted individual larval feeding galleries from each of the parental galleries. We recorded (1) mating success or failure, (2) parental gallery length (cm), and (3) total sum of eggs and larval offspring. Reproductive success was determined by the presence of eggs or larval galleries. In unsuccessful pairs, no vertical gallery was present, or the pair of adult beetles were dead with no evidence of eggs or larval feeding galleries.

### Diapause status of field-sampled beetles

The laboratory experiments described above tested whether adult spruce beetles could reproduce in constant warmth without cold temperature cues, but they do not reflect field-relevant conditions. To assess how field temperatures affect reproductive maturity, we collected overwintering adult beetles directly from infested trees at Guanella Pass, Colorado during fall 2019 and fall and spring 2021 and 2022. Initially, we collected beetles in fall to establish a time point where all beetles were clearly in adult reproductive diapause (Fig. 3). However, after our first collection we realized that most female beetles in the field were reproductively mature in the fall (see results). Thereafter, we made several more fall collections over the course of two collection years to confirm this observation, then followed with a spring collection to test whether egg chambers and oocytes might be flexibly re-absorbed as has been documented in other insects under unfavorable environmental conditions (Nozaki & Matsuura, 2021). This allowed us to confirm whether females with mature ovaries survive overwinter.

We identified up to 20 infested trees each field season (**Fig. S3**) and collected 51 and 72 female beetles in 2019 and 2021, respectively, over several collection dates in the fall, and 29 in the spring of 2022 (Table 1). We collected beetles overwintering at the base of standing, infested trees by carefully removing sections of bark, and all beetles found within the phloem were placed in a petri dish with moist filter paper. All samples were transported to the laboratory in a cooler with an ice pack, and subsequently frozen live for storage prior to dissections to assess ovary maturity.

**Table 1.** Number of mature female field-collected spruce beetle adults from the base of infested trees. Beetles were collected from Guanella Pass, Colorado before, during and after winter.

| Date Collected | Number Dissected | Number Mature |
| --- | --- | --- |
| 9/27/2019 | 22 | 17 |
| 10/4/2019 | 6 | 4 |
| 10/31/2019 | 7 | 7 |
| 11/20/2019 | 1 | 1 |
| 12/4/2019 | 15 | 15 |
| 10/6/2021 | 31 | 27 |
| 11/9/2021 | 41 | 35 |
| 4/15/2021 | 29 | 25 |

We also monitored field temperatures to assess whether transient exposure to cold temperatures interspersed with warmer temperatures favorable for development could explain the observation that female beetles had matured ovaries by late September. We were not able to install loggers in late summer of 2019. Instead, we installed them from 5 August 2023 to 14 October 2023, used data from two nearby SNOTEL stations (Jackwacker Gulch and Echo Lake) to parameterize a linear model, then projected 2019 phloem temperatures using the model. We removed a 5.1 x 7.6 cm section of phloem, inserted data loggers (Thermochron iButton DS 1922L-f5#) wrapped in parafilm and set to record temperatures in hourly increments (±1°C), then stapled the phloem flap back in place. We placed loggers in two trees in close proximity to the trees from which we field-collected beetles: one on Naylor Lake Road, and one slightly downhill at Guanella Pass Campground.

### Statistical Analysis

We estimated the 95% confidence intervals and performed hypothesis tests for all reported proportions (proportion females with mature ovaries and proportion of successful matings) using either the binom.test (exact binomial test), fisher.test (Fisher’s exact test), or glm (generalized linear model for logistic regression with family = ‘binomial’ with a logit link function) functions in R (R version 3.4.4). We fit a linear model (lm function) to predict 2023 field phloem daily maximum and minimum temperatures from 2023 SNOTEL data. We fit separate models for each tree with an installed logger with SNOTEL data from each station as continuous covariates, then used the fitted models to predict 2019 phloem temperatures using 2019 SNOTEL data from 5 August to 27 September (the first field beetle collection date in 2019). We constructed distributions of maximum and minimum temperatures over this time period based on the mean predicted temperatures of the two trees.

## Results

### Adults emerged without cold treatment

A total of 4208 adults emerged from Colorado-origin bolts and 4530 from Wyoming-origin bolts. Adult emergence from the Wyoming bolts began immediately after they were placed at 22°C (**Fig. 5a**), and they continued to emerge regularly for 211 days. One emergence peak was present for Wyoming during the week of 5 September 2018 (n=845; 55 days at 22°C).

**Figure 5:**
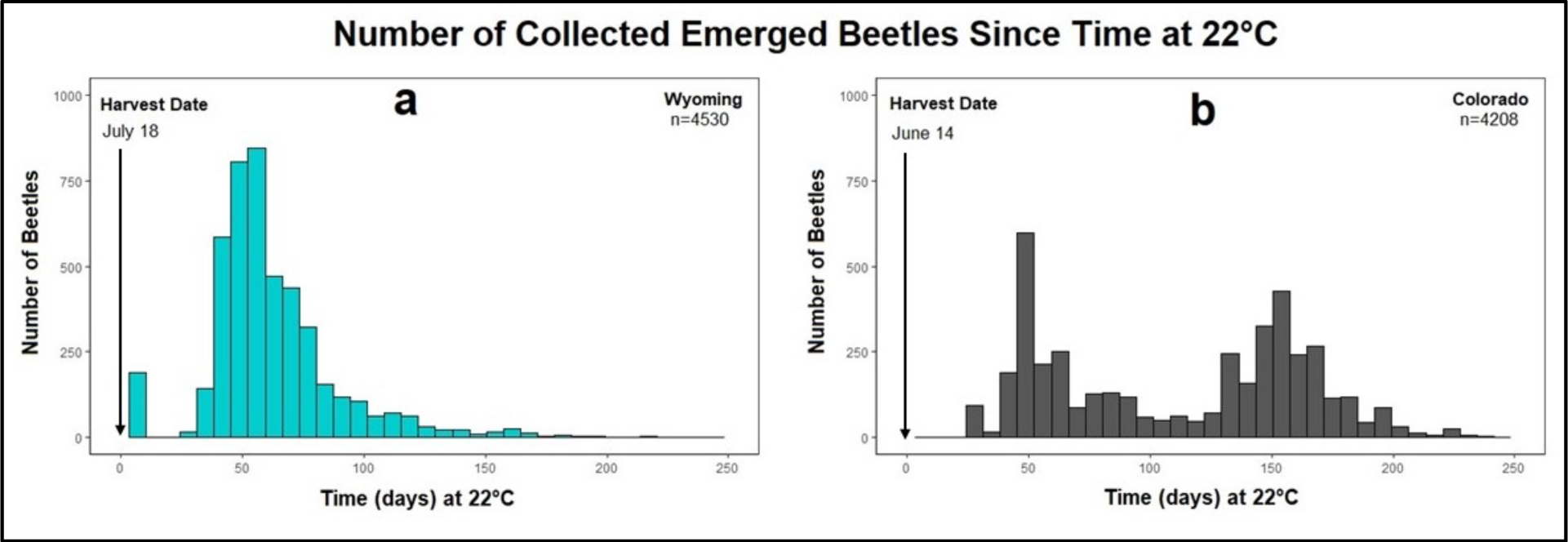
Number of emerged adult beetles that were collected following the day placed at 22°C (day 0) until 5 days passed with no newly emerged beetles (day 246) for (a) Wyoming and (b) Colorado populations.

Adult emergence from Colorado bolts began 25 days after they were placed at 22°C (**Fig. 5b**), and they continued to emerge regularly for 217 days. Two emergence peaks were present for Colorado, one during the week of 26 July 2018 (n=598; 48 days at 22°C) which was likely made up of re-emerged parents, and the other during the week of 8 November 2018 (n=427; 153 days at 22°C).

### Some, but few females were reproductively mature under constant warm conditions

Though beetles emerged from bolts at a high rate, most emerging female beetles from both the northern and southern collection sites were not reproductively mature. Given that these adults did not experience any chilling, this result suggests that most individuals had entered a reproductive diapause that had not been terminated by cold. A small proportion of adults from each population, however, developed mature ovaries in the constant warm temperatures. Of the adult females dissected, 11% of Colorado beetles (9 out of 83 total, 95% CI of proportion mature; 0.051 - 0.20) and 1% of Wyoming beetles (1 out of 94 total, 95% CI of proportion mature; 0.00027 - 0.058) were mature (**Fig. 6a**).

**Figure 6:**
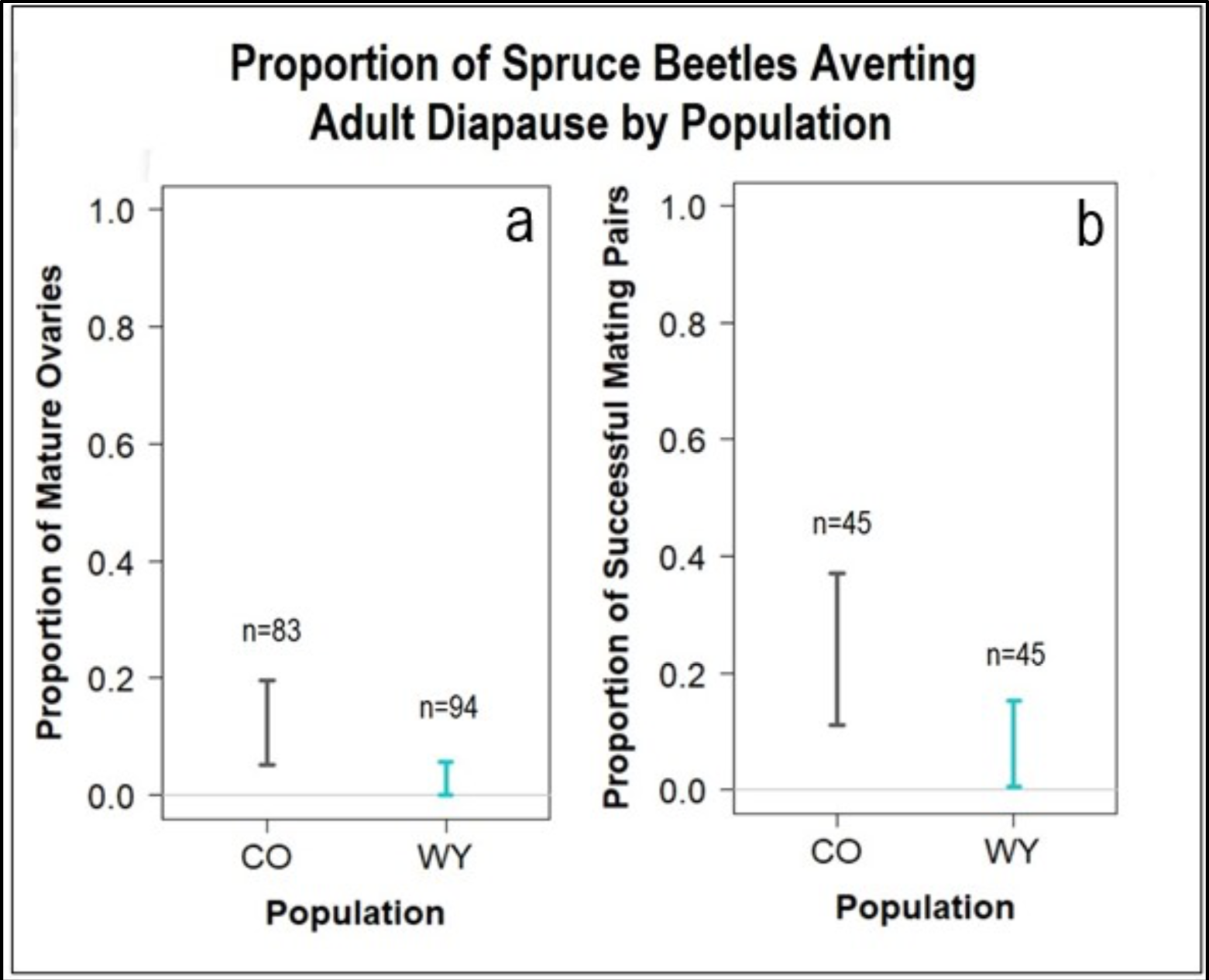
95% Confidence intervals for the proportion of spruce beetles averting adult diapause per population. Proportion of dissected females that were mature after 100 days at 22°C, evidenced by presence of eggs (a). Proportion of mating pairs that produced offspring after 50-70 days at 22°C (b). All proportions were greater than zero (p << 0.001) based on binomial tests of the null hypothesis of proportion mature/proportion mating = 0.

With these relatively large sample sizes, there is evidence that the proportion was higher for Colorado compared to Wyoming beetles (*P* = 0.007, Fisher’s exact test), though the 95% CIs do not provide compelling evidence for a substantial difference. Only adults that emerged after 100 days at a warm temperature were scored. Within the range of emergence times over which the samples were collected (100 – 246 days), emergence time had no discernable relationship to the probability that a female had mature ovaries based on a generalized linear model with a logit link function (CO – Wald Z = 0.012, p=0.522; WY – Wald Z = 0.025, p=0.344 F**ig. S4**).

### Some, but few adults successfully reproduced under constant warm conditions

Corroborating our observations of a low but non-zero rate of reproductive maturity based on ovarian development, some male-female pairs were able to successfully reproduce without a cold treatment. Of the 45 mating pairs from Colorado, 22% reproduced successfully (10 out of 45 pairs; 95% confidence interval of the proportion: 0.11 - 0.37), and 4% of Wyoming mating pairs successfully reproduced (2 out of 45 pairs; 95% confidence interval of the proportion: 0.0054 - 0.15) (**Fig. 6b**). As was the case for ovary maturity, the proportion of successfully reproducing females was higher in the Colorado population compared to the Wyoming population (*<u>P</u>* <u>= 0.03, Fisher’s exact test</u>), but only marginally (see 95% CI in Fig. 6b). Female size did not differ substantially between successful and unsuccessful reproducing females (**Fig. S5**).

### Field-collected adult beetles from Colorado are reproductively mature in early fall and spring

Most adult female beetles collected from the base of trees during late summer through early spring had mature ovaries, including on the first 2019 collection date of 27 September (Table 1). From 5 August to the first beetle collection date (27 September) in 2019 predicted daily minimum phloem temperatures occasionally dipped below 4°C, while predicted daily maximum temperatures frequently surpassed 15°C (Fig. 7). It appears that the combination of occasional cold temperature exposure with temperatures permissive for growth and morphogeneis was sufficient to terminate diapause and allow ovary maturation, in contrast to our observations of little to no ovary maturation in constant warm temperatures in the lab. The high percentage of spring-collected beetles with mature ovaries suggest that egg chambers and oocytes are likely not resorbed, and that beetles with mature ovaries do survive winter. We also observed that a random sample of females were mated prior to winter based on presence of sperm in the spermatheca (27 September 2019: n=3; 4 October 2019: n=1; 31 October 2019: n=2; 4 December 2019: n=5). This suggests that some female beetles mated prior to relocation to the base of trees for overwintering. Because we only collected beetles that overwintered in clusters at the base of trees that were infested the previous year, it is unlikely that our observations of mature ovaries prior to and after adult overwintering are misidentifications of beetles that had already overwintered one year as adults.

**Figure 7:**
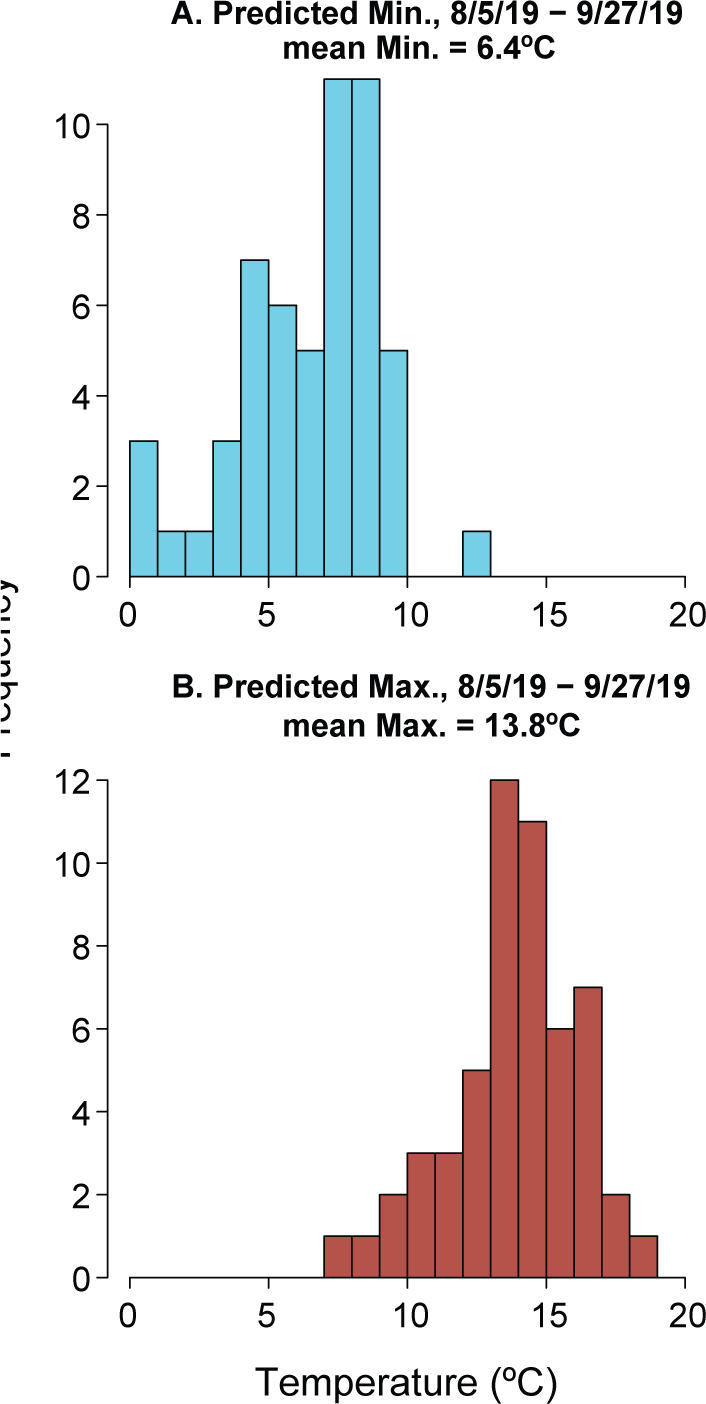
Distributions of predicted daily minimum and maximum field phloem temperatures between 5 August and 27 September 2019 at the Guanella pass site where adult beetles where collected in the field. Predictions were derived from linear models predicting 2023 field phloem temperature from nearby SNOTEL data as described in the methods. Distributions represent the mean daily min/max between the two trees with temperature loggers.

## Discussion

### Spruce beetle reproductive development is highly dependent on low temperature exposure

Reproductive diapause in the spruce beetle appears to be heavily influenced by exposure to low temperature. Previously, Bleiker and Willsey (2020) showed that pulses of cold temperatures can accelerate and synchronize beetle emergence from bolts in the lab, although reproductive maturity of these insects was not evaluated. We found that large numbers of beetles emerged from infested bolts kept in constant warmth (22°C), but only 11% and 1% of females, from the Colorado and Wyoming sites, respectively, had mature ovaries. Similarly, only a proportion of male-female pairs from each population reproduced successfully; 22% of Colorado beetles and 4% of Wyoming beetles. Collectively, these results suggest that a small proportion of spruce beetles did not enter what was presumed to be an obligate reproductive diapause, and the proportion of non-diapausing individuals was higher in the more southern Colorado population. This result confirms unpublished observations cited in Schebeck et al. (2017).

Moreover, without cold cues the extended emergence time observed in the lab (> 200 days) differed substantially from the more synchronous emergence that occurs within ∼30 days in the field (McCambridge & Knight, 1972). This synchronized, mass emergence, thought to be facilitated by adult reproductive diapause, is ecologically necessary for spruce beetles to effectively overcome host tree defenses (Chippendale, 1982; Raffa *et al*., 2008). In the case of the field-collected Colorado beetles with mature ovaries as early as late September, we suspect that temperatures below the flight threshold of near 13 °C (Holsten & Hard, 2001) prevented dispersal and attacks on new hosts. That is, reproductive diapause delayed dispersal during an unfavorable time of year (i.e., August and September) until simple temperature regulation was adequate to ensure synchronized emergence the following spring.

In contrast to lab rearing in constant warmth, a majority of spruce beetles sampled from infested trees in the field during fall and early winter were reproductively mature, an unexpected result. Relatively short exposures to transiently (i.e., diurnally) cold temperatures appeared to be sufficient to potentiate reproductive development, consistent with a body of literature illustrating major differences in insect life history in fluctuating versus constant temperatures (Colinet *et al*., 2015). Although we do not know when during spring or summer 2019 the adult life stage was attained at our study locations, we estimate that the adults collected in mid-September experienced minimum temperatures less than or equal to 2°C on at least 4 days. Constant exposure to 2°C for 25 days in the lab stimulated diapause termination in *D. rufipennis* (Bleiker & Willsey, 2020), and our data suggest that much shorter, though frequent pulses of comparable or even more moderate cold may be sufficient for termination. Insects with a reproductive diapause are thought to overwinter in reproductive arrest, delaying gamete maturation until conditions are more permissive in spring or summer (Hodek, 2011). It is possible that reproductive diapause in spruce beetle may primarily act as a block to prevent reproduction prior to winter, rather than serving as the cold-hardy, environmentally buffered stage that we typically attribute to a winter diapause. Females that had already started to invest in reproductive development were clearly capable of withstanding winter cold and nutrient reserve depletion to successfully reproduce the following year. Initiating reproductive development prior to winter may facilitate early establishment of new broods the following spring.

### Potential responses to changing climates

Our findings that a small proportion of adult spruce beetles can become reproductively mature without cold temperature cues, in addition to reproductive development prior the onset of continuous cold during winter, highlight further flexibility in the spruce beetle lifecycle. In addition to a facultative prepupal diapause, a facultative reproductive diapause in some individuals could allow completion of a generation in a single summer, like that observed for mountain pine beetle at a warm location in southern California when overwintered parents re-emerged in June and attacked new trees (Bentz *et al*., 2014). Moreover, warm daytime temperatures with a few pulses of cold exposure through transient cold fronts or accumulated cold nights could in combination enable both reproductive maturation and successful mating and reproduction in late summer or fall. The potential for two generations in a year, however, would be constrained by cold winter temperatures that would reduce development time and potentially induce the prepupal diapause, as was found for mountain pine beetle (Bentz & Powell, 2014).

Autumn and winter temperatures would have to warm catastrophically and be consistently above 15°C to avoid a prepupal diapause, conditions in spruce habitats that are not forecasted by Global Climate Models (Bentz *et al*., 2022). Mating and reproduction in the fall could also result in life stages not adapted for cold (i.e., eggs, early instars). Flexibility of adult reproductive development could provide alternate pathways between semivoltinism and univoltinism, but could also lead to fractional voltinism that could have positive or negative influences on population dynamics. Further studies employing shorter chilling periods in the lab and earlier, more fine-grained field sampling would further elucidate the flexibility of adult reproductive diapause in nature.

Variation in the adult reproductive diapause response may also provide a pathway to evolutionary changes in the life cycle. Indeed, evidence for absolute chilling thresholds for diapause termination in insects is relatively scarce. Rather, diapausing insects that are temperature sensitive often exhibit variable responses within and across populations (Masaki, 1961; Feder *et al*., 1997; Schmidt *et al*., 2005), where cold temperatures accelerate diapause development, but are not absolutely necessary for diapause termination (Hodek, 2013). And, diapause regulation, and subsequently field phenology, can evolve extremely rapidly in insects (Gomi *et al*., 2007; Bradshaw & Holzapfel, 2008; Batz *et al*., 2020). Though the proportion was small, some adult beetles from both Colorado and Wyoming matured ovaries and successfully reproduced without any low temperature exposure. If this variation were heritable, warming conditions could select for these currently rare phenotypes, moving the population mean response towards earlier reproductive maturity. Of course, warming conditions could also plausibly select against the warm-reproducing phenotype, e.g. if fractional voltinism leads to overwintering of life stages that cannot survive sustained and more extreme winter cold. Better understanding of these possibilities would be challenging to achieve, likely requiring both field observation and lab manipulation of temperature and tracking of phenotypic (and potentially genotypic) distributions across generations.

### Reproductive maturity, not emergence, is the most reliable indicator of adult diapause status

Previous studies have used emergence as a measure of adult diapause termination, including those on spruce beetle (Bleiker & Willsey, 2020) and the Douglas-fir beetle, *Dendroctonus pseudotsugae* (Ryan, 1959). These studies correlate activity outside of the tree (e.g., being capable of flight) with reproductive maturity. However, our study shows that most emerged spruce beetles are not reproductively mature in the absence of a cold treatment. Because a high number of adults emerged from both populations but only a small proportion were reproductively viable, these results also suggest that adult emergence does not necessarily indicate reproductive maturity. Immature adults of this species are capable of activity – some proportion relocate from the upper bole to the base of the tree prior to overwintering, one year before they would normally reproduce (Knight, 1961). In combination with our results, this suggests that emergence under constant warm conditions in the lab may primarily reflect relocation behavior, not reproductive maturity.

We scored reproductive maturity using both ovary dissections and mating trials and found that the proportion of mating success without a cold treatment was higher than the proportion of adults with mature ovaries for both Colorado and Wyoming populations. This could be due to several factors including changes in behavior when exposed to a potential mate or a new uninfested tree (i.e., green rearing bolts). Individual beetle variation may be another consideration as beetles used for ovary maturity assessments were sampled across the emergence distribution after 100 days at 22°C, whereas beetles used from mating experiments were sampled at the beginning of the main emergence curves, after approximately 50-70 days at 22°C. Thus, the experimental conditions and timing of sampling may influence estimates of the proportion of diapausing vs. mature bark beetle adults in a population.

### Summary

Our study documents a low, but measurable rate of successful adult reproduction without chilling, and geographic variation in the thermal sensitivity of adult reproductive diapause in spruce beetle: beetles from a southern site had greater propensity to avert reproductive diapause than beetles from a northern site. Emergence was a poor predictor of reproductive maturity under warm conditions. We further showed that adult female beetles can and usually do fully mature ovaries prior to winter. Thus, field observations of overwintering developmental status differ from predictions based on laboratory studies and conventional wisdom about adult reproductive diapause. Further investigation of the thermal sensitivity of development and variation within populations could lead to better predictions about how beetle populations may respond to changing seasonal temperatures under climate change.

## ACKNOWLEDGEMENTS

We wish to thank Jim Vandygriff, Alex Rudney, Brian Verhulst, Taylor Siddons, and Isaac Dell for help with field beetle collections, Sky Stephens, Rebecca Stokes, Mac Calvert, Jantina Toxopeus, Joshua Gordon, Nooshin Sanaei, Matin Sanaei, Manaal Dalwadi, Monica Khazaal, Isaiah Sower, Lahari Gadey, and Colin McSweeneyfor help with beetle rearing and ovary dissections, Erika Eidson and Kristy Branch for help with figure design, and Corinne Casper and Andrew Neisess for assistance placing and retrieving field temperature loggers. This work was supported by NSF DEB 1638951 to GJR, and in part by the U.S. Department of Agriculture, Forest Service, Rocky Mountain Research Station. The findings and conclusions in this publication are those of the authors and should not be construed to represent any official USDA or U.S. Government determination or policy.

## DATA AVAILABILITY

All data will be provided at the time of publication in a peer-reviewed journal.

---

**Figure S1:**
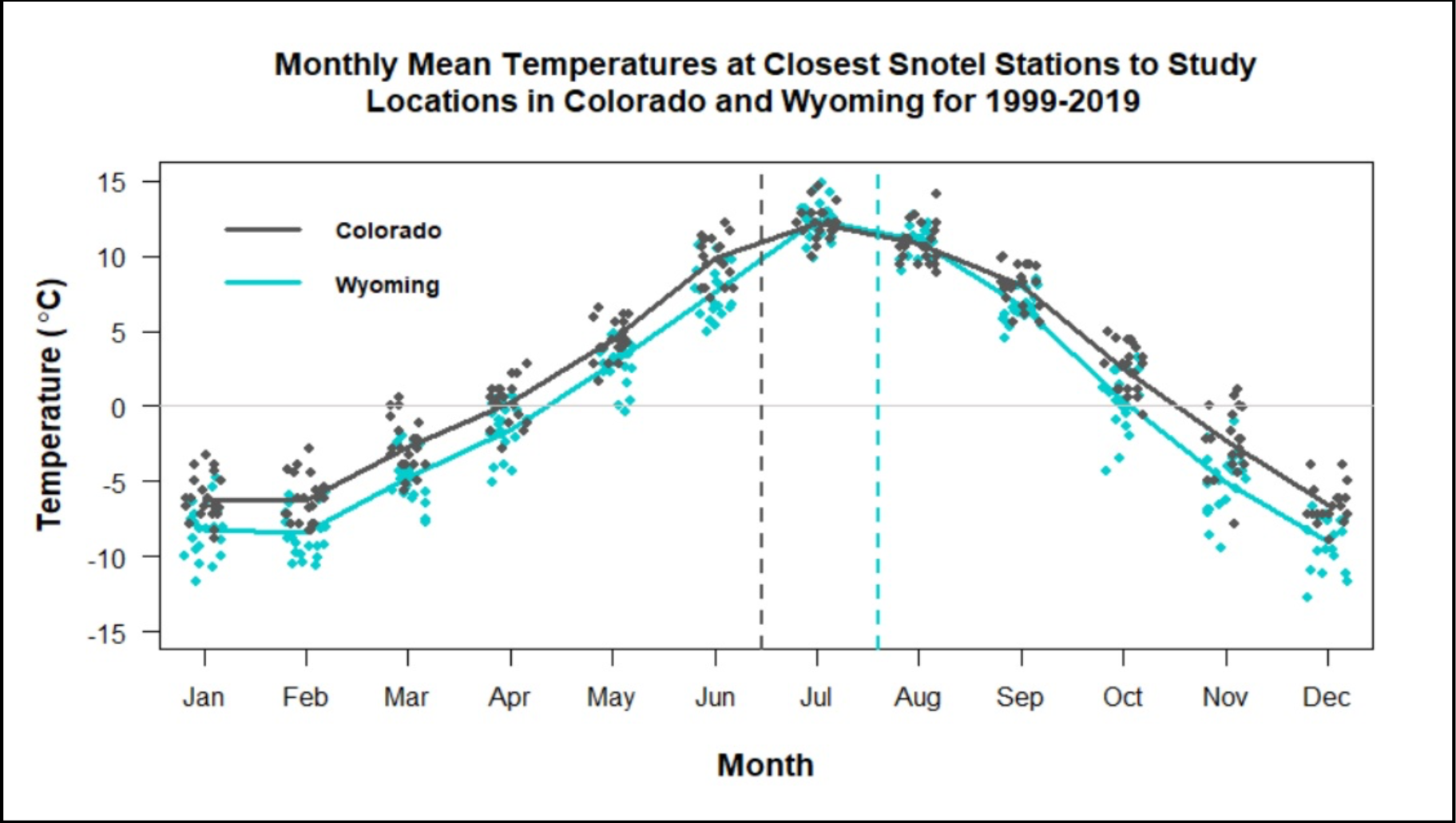
Monthly mean temperatures for closest weather stations (https://www.ncdc.noaa.gov/cdo-web/) to each study location in Colorado and Wyoming for 1999-2019. Dates infested trees were harvested are shown in vertical dashed lines. Colorado was cut on June 14, 2018 and Wyoming on July 18, 2018.

**Figure S2:**
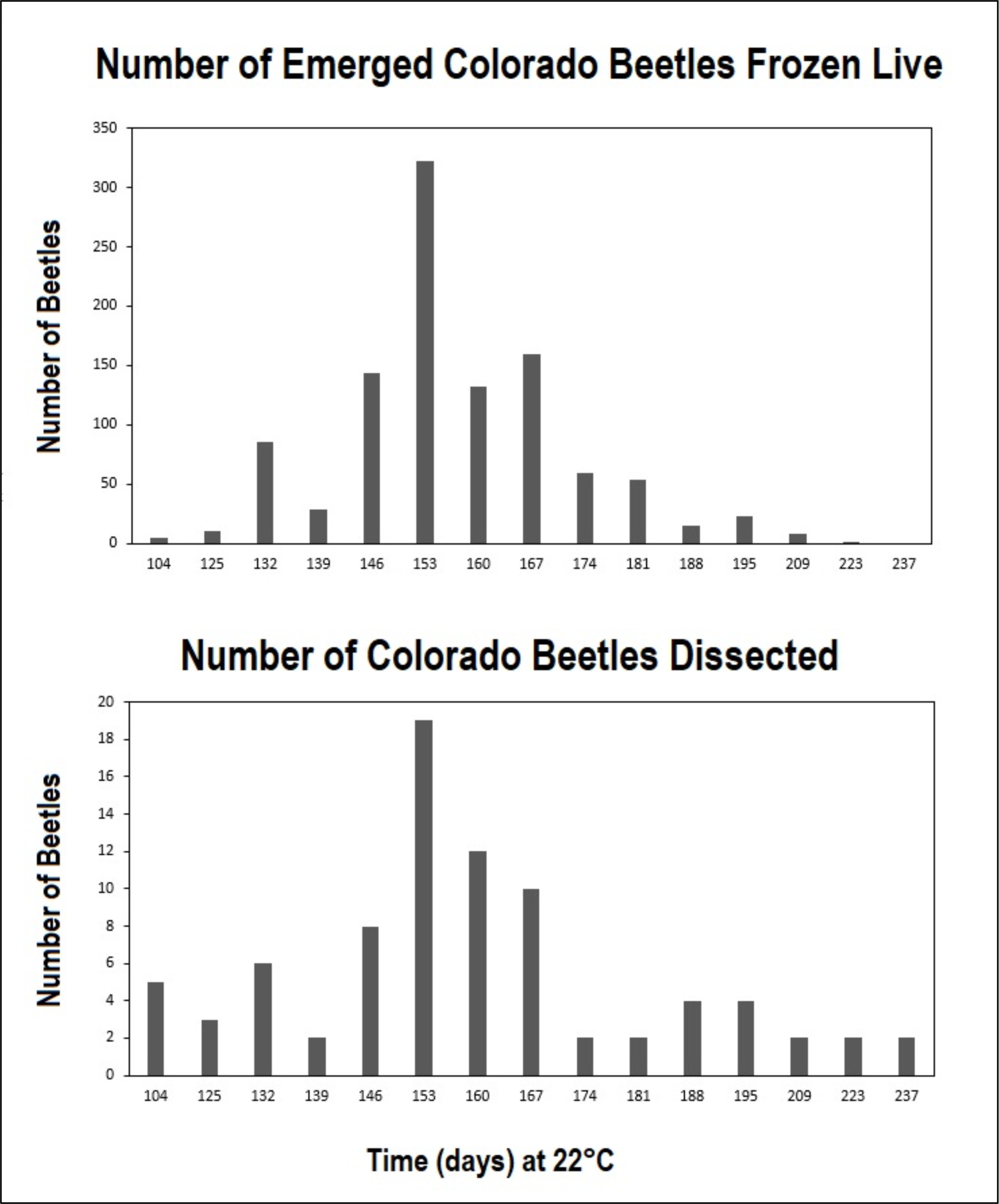
*Top:* Distribution of live frozen emerged Colorado beetles from which beetles were sampled for ovary dissection. *Bottom:* Sampling distribution of number of beetles at time at 22°C.

**Figure S3:**
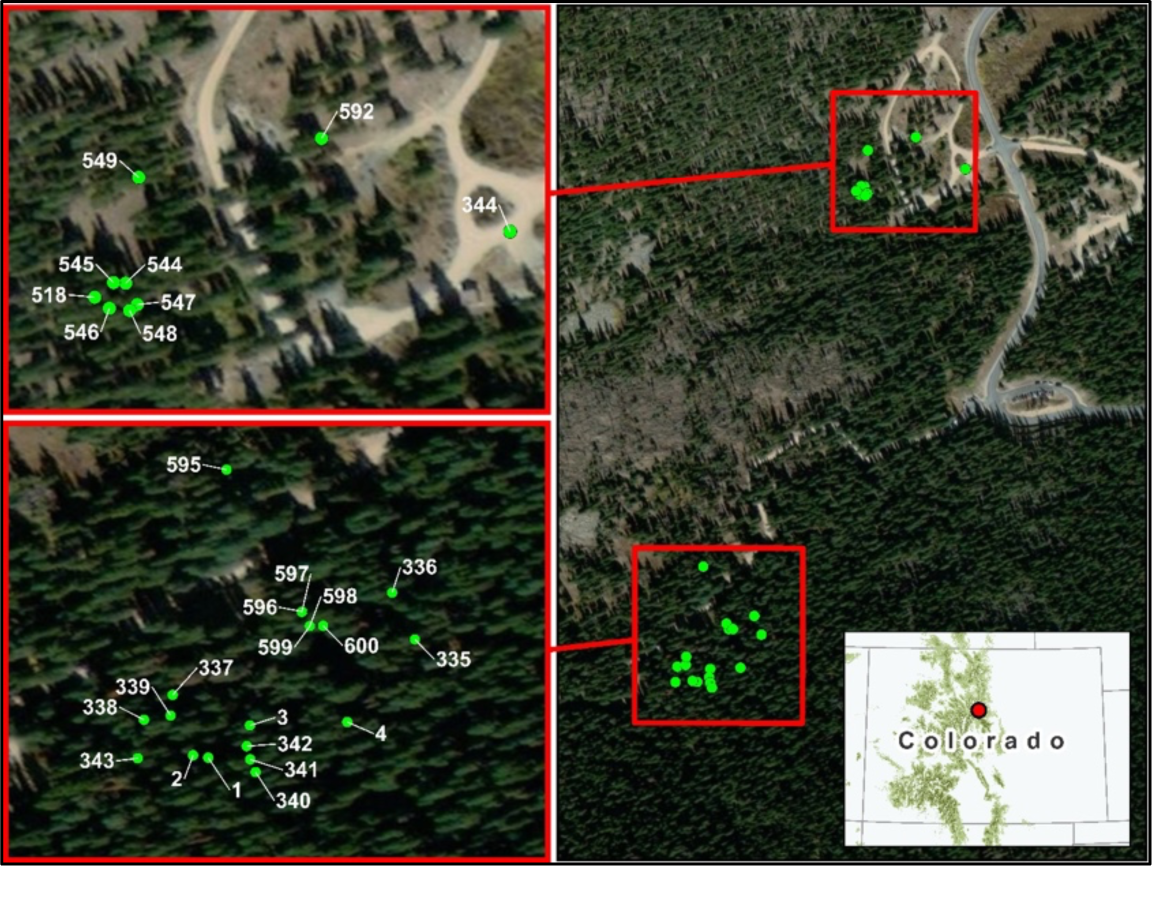
Map showing infested trees located in Guanella Pass Campground (*top*) and off Naylor Lake Road (*bottom*) in Colorado that were flagged early Fall 2019 for later winter field collection of beetles overwintering at the base.

**Figure S4:**
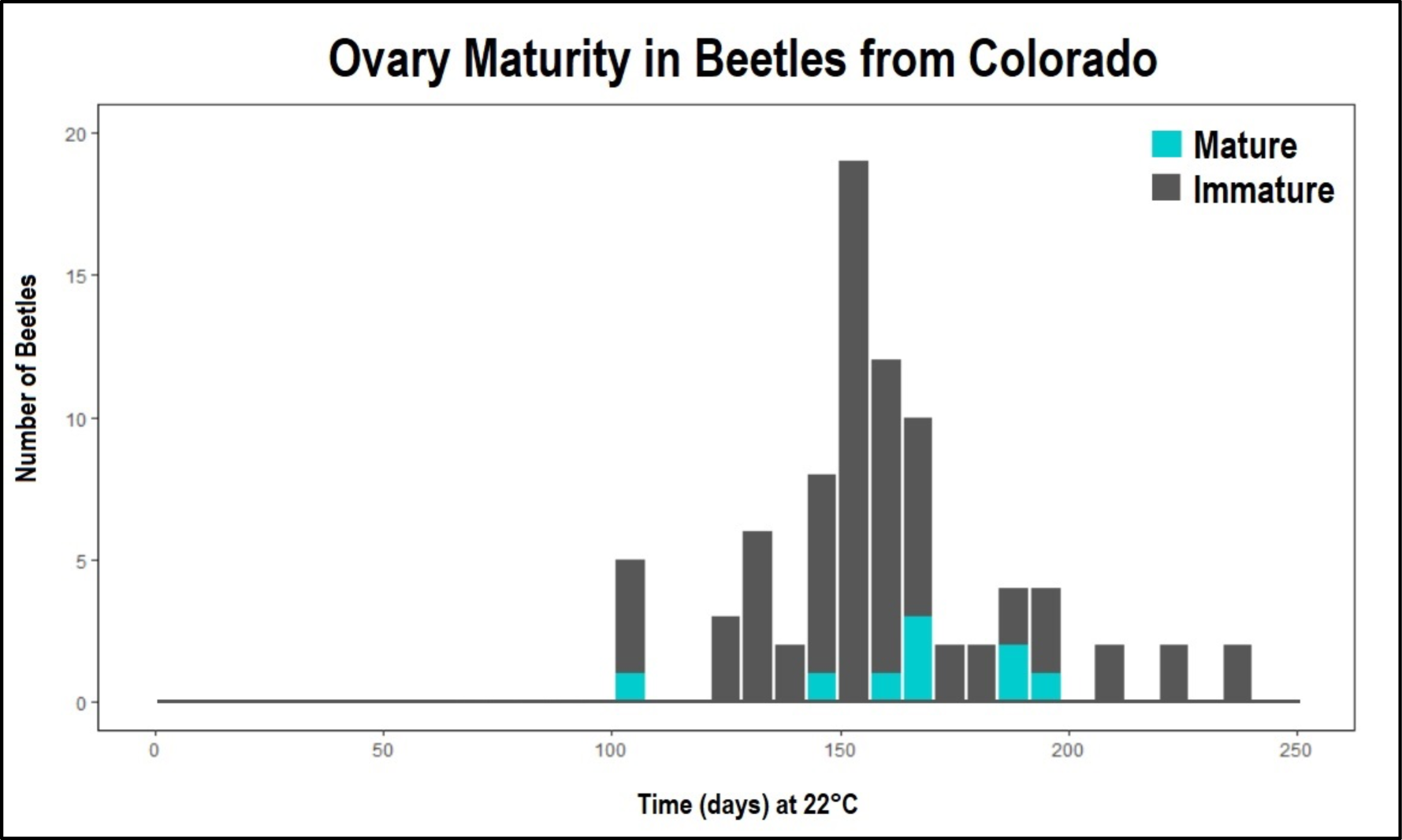
Number of emerged adult spruce beetles with immature and mature ovaries from the Colorado population of infested bolts kept at 22°C following harvest. Adults that emerged prior to day 100 following being placed at 22°C were not sampled or dissected.

**Figure S5:**
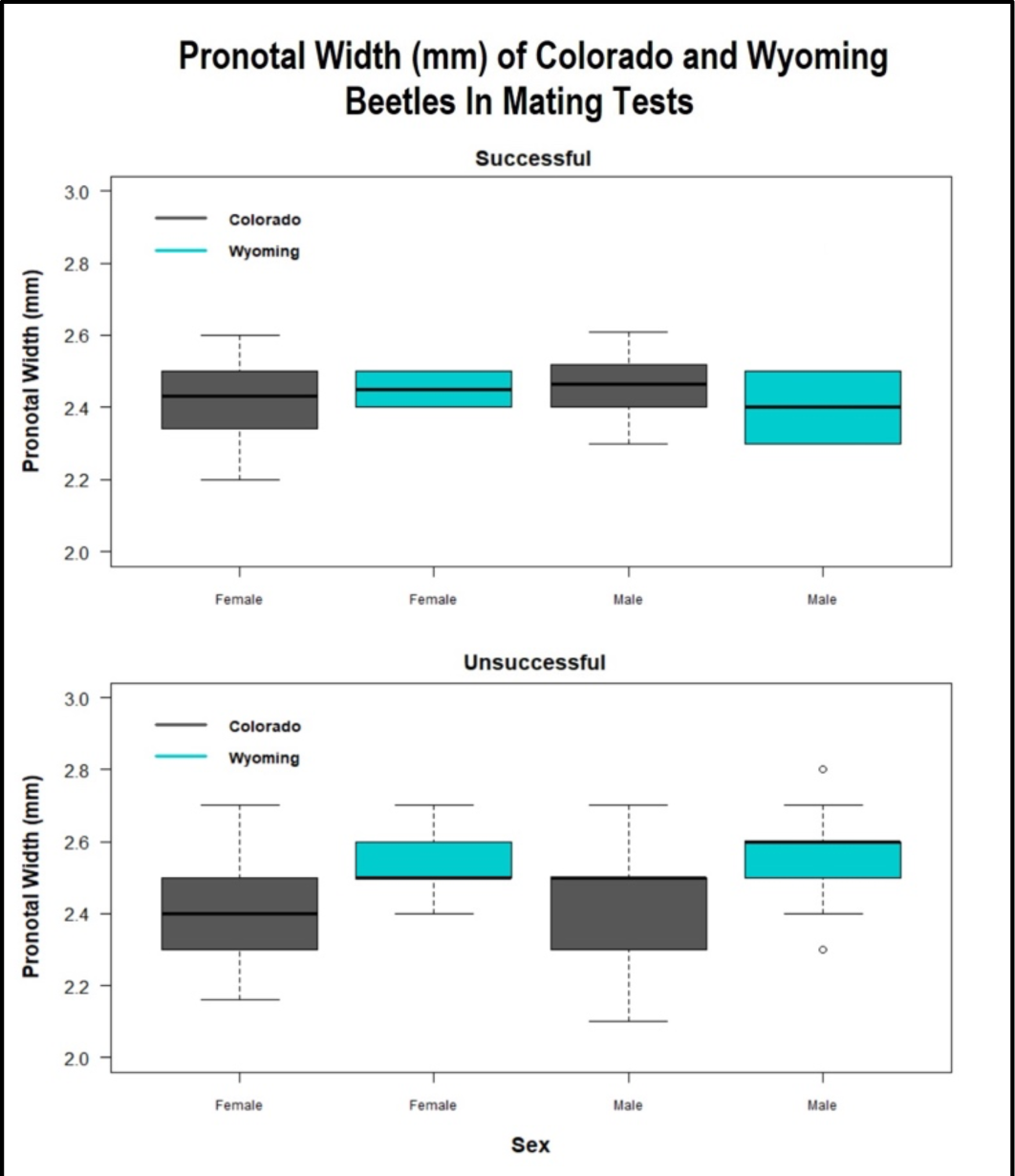
Pronotal width of successful mating pairs by sex (*top)* and unsuccessful pairs (*bottom*) for Colorado and Wyoming beetle populations.

**Table S1:** Trees collected during summer of 2018 for laboratory experiments. One infested tree was cut in Colorado and the other in Wyoming. A third healthy and uninfested tree was harvested in Colorado to be used in mating trials.

| Status | Location | Date Cut | Infestation Year | Elevation (m) | dbh (cm) | Coordinates | County |
| --- | --- | --- | --- | --- | --- | --- | --- |
| Infested (Southern) | Guanella Pass | 6/14/2018 | 2017 | 3478.99 | 43.69 | N 39.60663°<br>W 105.72268° | Clear Creek, CO |
| Infested (Northern) | Togwotee Pass | 7/18/2018 | 2017 | 2595.98 | 29.72 | N 43.81040°<br>W 110.21263° | Teton, WY |
| Green | Turquoise Lake | 8/13/2018 | NA | 3310.13 | 29.72 | N 39.264200°<br>W 106.436517° | Lake, CO |

## REFERENCES

Altermatt, F. (2010) Tell me what you eat and I’ll tell you when you fly: diet can predict phenological changes in response to climate change. Ecology Letters, 13, 1475–1484.

Batz, Z.A., Clemento, A.J., Fritzenwanker, J., Ring, T.J., Garza, J.C. & Armbruster, P.A. (2020) Rapid adaptive evolution of the diapause program during range expansion of an invasive mosquito. Evolution, 74, 1451–1465.

Bentz, B., Vandygriff, J., Jensen, C., Coleman, T., Maloney, P., Smith, S., et al. (2014) Mountain Pine Beetle Voltinism and Life History Characteristics across Latitudinal and Elevational Gradients in the Western United States. Forest Science, 60, 434–449.

Bentz, B.J., Hansen, E.M., Davenport, M. & Soderberg, D. (2022) 2 - Complexities in predicting mountain pine beetle and spruce beetle response to climate change. In Bark Beetle Management, Ecology, and Climate Change (ed. by Gandhi, K.J.K. & Hofstetter, R.W.). Academic Press, pp. 31–54.

Bentz, B.J., Jönsson, A.M., Schroeder, M., Weed, A., Wilcke, R.A.I. & Larsson, K. (2019) Ips typographus and Dendroctonus ponderosae Models Project Thermal Suitability for Intra-and Inter-Continental Establishment in a Changing Climate. Frontiers in Forests and Global Change, 2.

Bentz, B.J. & Powell, J.A. (2014) Mountain Pine Beetle Seasonal Timing and Constraints to Bivoltinism: A Comment on Mitton and Ferrenberg, “Mountain Pine Beetle Develops an Unprecedented Summer Generation in Response to Climate Warming.” The American Naturalist, 184, 787–796.

Berg, E.E., David Henry, J., Fastie, C.L., De Volder, A.D. & Matsuoka, S.M. (2006) Spruce beetle outbreaks on the Kenai Peninsula, Alaska, and Kluane National Park and Reserve, Yukon Territory: Relationship to summer temperatures and regional differences in disturbance regimes. Forest Ecology and Management, Spruce beetles and forest ecosystems of south-central Alaska, 227, 219–232.

Bleiker, K.P. & Willsey, T. (2020) Experimental Evidence Supporting an Obligate Adult Diapause for Spruce Beetle (Coleoptera: Curculionidae) from British Columbia. Environmental Entomology, 49, 98–103.

Bradshaw, W.E. & Holzapfel, C.M. (2008) Genetic response to rapid climate change: it’s seasonal timing that matters. Molecular Ecology, 17, 157–166.

Buckley, L.B., Nufio, C.R., Kirk, E.M. & Kingsolver, J.G. (2015) Elevational differences in developmental plasticity determine phenological responses of grasshoppers to recent climate warming. Proceedings of the Royal Society B: Biological Sciences, 282, 20150441.

Chippendale, G.M. (1982) Insect Diapause, the Seasonal Synchronization of Life Cycles, and Management Strategies. Entomologia Experimentalis et Applicata, 31, 24–35.

Colinet, H., Sinclair, B.J., Vernon, P. & Renault, D. (2015) Insects in Fluctuating Thermal Environments. Annual Review of Entomology, 60, 123–140.

Dambroski, H.R. & Feder, J.L. (2007) Host plant and latitude-related diapause variation in Rhagoletis pomonella: a test for multifaceted life history adaptation on different stages of diapause development. Journal of Evolutionary Biology, 20, 2101–2112.

Denlinger, D.L. (2022) Insect diapause. Cambridge University Press.

Deutsch, C.A., Tewksbury, J.J., Huey, R.B., Sheldon, K.S., Ghalambor, C.K., Haak, D.C., et al. (2008) Impacts of climate warming on terrestrial ectotherms across latitude. Proceedings of the National Academy of Sciences, 105, 6668–6672.

Dyer, E.D.A. & Hall, P.M. (1977) FACTORS AFFECTING LARVAL DIAPAUSE IN DENDROCTONUS RUFIPENNIS (COLEOPTERA: SCOLYTIDAE). The Canadian Entomologist, 109, 1485–1490.

Eidson, E.L., Mock, K.E. & Bentz, B.J. (2018) Low offspring survival in mountain pine beetle infesting the resistant Great Basin bristlecone pine supports the preference-performance hypothesis. PLOS ONE, 13, e0196732.

Feder, J.L., Stolz, U., Lewis, K.M., Perry, W., Roethele, J.B. & Rogers, A. (1997) The Effects of Winter Length on the Genetics of Apple and Hawthorn Races of Rhagoletis Pomonella (diptera: Tephritidae). Evolution, 51, 1862–1876.

Forrest, J.R. (2016) Complex responses of insect phenology to climate change. Current Opinion in Insect Science, Global change biology * Molecular physiology, 17, 49–54.

Forrest, J.R.K. & Thomson, J.D. (2011) An examination of synchrony between insect emergence and flowering in Rocky Mountain meadows. Ecological Monographs, 81, 469–491.

Gomi, T., Nagasaka, M., Fukuda, T. & Hagihara, H. (2007) Shifting of the life cycle and life-history traits of the fall webworm in relation to climate change. Entomologia Experimentalis et Applicata, 125, 179–184.

Hansen, E.M. & Bentz, B.J. (2003) Comparison of reproductive capacity among univoltine, semivoltine, and re-emerged parent spruce beetles (Coleoptera: Scolytidae). The Canadian Entomologist, 135, 697–712.

Hansen, E.M., Bentz, B.J., Powell, J.A., Gray, D.R. & Vandygriff, J.C. (2011) Prepupal diapause and instar IV developmental rates of the spruce beetle, Dendroctonus rufipennis (Coleoptera: Curculionidae, Scolytinae). Journal of Insect Physiology, 57, 1347–1357.

Hansen, E.M., Bentz, B.J. & Turner, D.L. (2001) Physiological basis for flexible voltinism in the spruce beetle (Coleoptera: Scolytidae). The Canadian Entomologist, 133, 805–817.

Hodek, I. (2011) Adult Diapause in Coleoptera. Psyche: A Journal of Entomology, 2012, e249081.

Hodek, I. (2013) Controversial aspects of diapause development. EJE, 99, 163–173.

Holsten, E.H. (1999) The Spruce Beetle. USDA Forest Service Forest Insect and Disease Leaflet No.27.

Holsten, E.H. & Hard, J.S. (2001) Dispersal flight and attack of the spruce beetle, Dendroctonus rufipennis, in south-central Alaska. U.S.D.A. Forest Service Research Paper PNW-RP-536.

Jacques, J., Sampaio, F., Santos, H.T. dos & Marchioro, C.A. (2019) Climate change and voltinism of Mythimna sequax: the location and choice of phenological models matter. Agricultural and Forest Entomology, 21, 431–444.

Knight, F.B. (1961) Variations in the Life History of the Engelmann Spruce Beetle. Annals of the Entomological Society of America, 54, 209–214.

Lyon, R.L. (1958) A Useful Secondary Sex Character in Dendroctonus Bark Beetles. The Canadian Entomologist, 90, 582–584.

Masaki, S. (1961) Geographic variation of diapause in insects. Bull. Fac. Agric. Hirosaki Univ., 7, 66–98.

Masaki, S. (2002) Ecophysiological consequences of variability in diapause intensity. European Journal of Entomology, 99, 143–154.

Massey, C.L. & Wygant, N.D. (1954) Biology and Control of the Engelmann Spruce Beetle in Colorado. U.S. Department of Agriculture.

McCambridge, W.F. & Knight, F.B. (1972) Factors Affecting Spruce Beetles during a Small Outbreak. Ecology, 53, 830–839.

McKee, F.R. & Aukema, B.H. (2015) Successful reproduction by the eastern larch beetle (Coleoptera: Curculionidae) in the absence of an overwintering period. The Canadian Entomologist, 147, 602–610.

Nozaki, T. & Matsuura, K. (2021) Oocyte resorption in termite queens: Seasonal dynamics and controlling factors. Journal of Insect Physiology, 131, 104242.

Nunes, M.V. & Saunders, D.S. (1989) The effect of larval temperature and photoperiod on the incidence of larval diapause in the blowfly, Calliphora vicina. Physiological Entomology, 14, 471–474.

Raffa, K.F., Aukema, B.H., Bentz, B.J., Carroll, A.L., Hicke, J.A., Turner, M.G., et al. (2008) Cross-scale Drivers of Natural Disturbances Prone to Anthropogenic Amplification: The Dynamics of Bark Beetle Eruptions. BioScience, 58, 501–517.

Ragland, G.J., Armbruster, P.A. & Meuti, M.E. (2019) Evolutionary and functional genetics of insect diapause: a call for greater integration. Current Opinion in Insect Science, Neuroscience • Special section on Evolutionary Genetics and Genomics, 36, 74–81.

Ryan, R.B. (1959) Termination of Diapause in the Douglas-Fir Beetle,. The Canadian Entomologist, 91, 520–525.

Safranyik, L., Simmons, C. & Barclay, H.J. (1990) A conceptual model of spruce beetle population dynamics. Information Report - Pacific and Yukon Region, Forestry Canada.

Schebeck, M., Dobart, N., Ragland, G.J., Schopf, A. & Stauffer, C. (2022) Facultative and obligate diapause phenotypes in populations of the European spruce bark beetle Ips typographus. Journal of Pest Science, 95, 889–899.

Schebeck, M., Hansen, E.M., Schopf, A., Ragland, G.J., Stauffer, C. & Bentz, B.J. (2017) Diapause and overwintering of two spruce bark beetle species. Physiological Entomology, 42, 200–210.

Schmidt, P.S., Matzkin, L., Ippolito, M. & Eanes, W.F. (2005) Geographic Variation in Diapause Incidence, Life-History Traits, and Climatic Adaptation in Drosophila Melanogaster. Evolution, 59, 1721–1732.

Tauber, M.J., Tauber, C.A. & Masaki, S. (1986) Seasonal adaptations of insects. Oxford University Press, USA.

Tobin, P.C., Nagarkatti, S., Loeb, G. & Saunders, M.C. (2008) Historical and projected interactions between climate change and insect voltinism in a multivoltine species. Global Change Biology, 14, 951–957.

Visser, M.E. & Both, C. (2005) Shifts in phenology due to global climate change: the need for a yardstick. Proceedings of the Royal Society B: Biological Sciences, 272, 2561–2569.

Werner, R.A., Holsten, E.H., Matsuoka, S.M. & Burnside, R.E. (2006) Spruce beetles and forest ecosystems in south-central Alaska: A review of 30 years of research. Forest Ecology and Management, Spruce beetles and forest ecosystems of south-central Alaska, 227, 195–206.

